# *In vitro* assessment of elevated soil iron on germinability and germination characteristics of *Sorghum bicolor* (L.) Moench after chemo-priming

**DOI:** 10.1101/2021.11.22.469542

**Authors:** Beckley Ikhajiagbe, Owalum Linus Onawo

## Abstract

The commercial importance of Sorghum (*Sorghum bicolor* (L.) Moench) has attracted breeders to increase its seed yield using various breeding approach. Adverse soil factors however hampered progress made in crop development, especially micronutrient toxicity. Plant growth stimulators (PGS) have a significant role in enhancing growth parameters in Sorghum. In the present study, seeds were primed in 50, 150, and 250 ppm of each of gibberellic acid, indole acetic acid, and ascorbic acid respectively for 1 hr before sowing in Petri dishesmoistened with 10 ml of the iron-rich solution obtained as filtrate from a mix of distilled water and ferruginous soil (1:1 v/w). Results showed that although germination percentage in ferruginous medium was significantly reduced, there was enhancement in germination percentagewhen the seeds were primed in gibberellic acid (GA). Germinability in the iron-rich medium was 31.2 hrs; this was significantly reduced to 19.6 to 21.1hrs when these seeds were primed with growth stimulators.Although, shoot length was significantly reduced in plants exposed to ferruginous solutions, the root parameters were however enhanced. They were no significant changes in the total number of root branches regardless of ferrugenic status or use of growth stimulating agents. The utilization of growth stimulators as priming agents is called for to reduce stress impacts imposed by ferruginous soils during germination.

## INTRODUCTION

Sorghum (*Sorghumbicolor*(L.) Moench) is one of the major staple crops for the world’s poorest and most food insecure people. Sorghum is one of the leading drought-tolerant crops, and is thus used as risk-preventive and strategic crop in numerous developing countries to address household food security. It is one of the primary sources of energy, protein, vitamins, and minerals for multitude of the poorest people in semi-arid areas (Stefoska-Needham *et al*.,2015). Sorghum is one of the mostparamount cereals in West Africa. It is mostly grown in semi-arid regionof tropics and subtropics, and in most West Africa countries, it suffices for 50% of total cereal crop land area.

Soil is a crucial part of successful agriculture and it’s the primary source of the nutrient that is used in raisingcrops (Valeraa, 1977). It consists of minerals organic matter, microbes, water and air (Nnadi *et al*., 2019). These constituentsof soil immensely affectthe fertility, buildup and porosity of diverse soils and as well impact the plant distribution (Ikhajiagbe et al., 2019). Inadequate soil nitrogen, limited phosphorus and iron toxicity are regarded as primary hindrances to the growth of several agricultural products (Adnan *et al*., 2018; Joshis et al., 2007). Similar soil characteristics have been related with red soils of the humid tropics and have always acted as indicators of severe weathering considering the wellspring of the parental materials as iron-rich mafic rock. These soils are also referred toas ferruginous ultisols (Cho and Ponnamperuma, 1971).Ferruginous ultisol are acidic red soils found in warm temperate, humid climates and in areas covered with deciduous or mixed forest (Yu *et al*., 2016). The total area of red soils is approximately 64 million km^2^, accounting for 45.2% of the earth’s surface area (Anumalla *et al*.,2019). In Nigeria, it is mainly seen in some southern state such as Edo state, occupying about seven zones, including extreme north and central Benin (Doyou *et al*., 2017). Humid climate tends to have high instances of red soil rich iron. The high iron in ferruginous soils form complex with soil insoluble phosphates hence reducing the bio-availability phosphorus to plants (Gyaneshwar *et al*., 2002). Red soil also has low amount of phosphorus in soils solution and ensues in frequent phosphorus deficiency of plants (Wang *et al*., 2014).

Yu *et al. (*2016), relates that red soil is generally derived from crystalline rock. There are usually malnourished growing soils with limited plant yield which affect productivity and food security.

Soil heavy metal concentrations may not however be totally due to industrial activities as some soils are originally ferruginous and therefore have increasingly high quantities of iron yet some others have increased levels of aluminium which predisposes such soils to more soil acidity (Ikhajiagbe,2016). Plant growth regulation (PGRs) or stimulators are chemicals which changes the plant growth and are applied in minute amounts. The germination and developmental mechanisms in plants are coordinated and been by the action and balance of separate group’s regulators, which may act as promoters or inhibitors of these processes. Many studies have shown that application of growth regulators enhance plant growth and crop yield (Hernandel, 1997). Plant growth stimulators has immense effects on plant growth and development, the exact action depending on the concentrations of the substances present and the response of the organs concerned. Applied gibberellic acid and indole-acetic acid showed increases in growth conditions (germination percentage, plant height, number of branches and leaves, total chlorophyll content, dry weight) (Gehan and Mona, 2011).

The fundamental priority for optimizing field production of any crop plant isthat it stands establishment. At suboptimal conditions the usual phenomenon include subsequent poor field establishment, poor seed germination. One of the mainhinderances to high yield and crop plant productionis the absence of synchronized crop establishment as a result of poor weather and soil conditions (Mwale *et al*.,2003).

Poor and unsynchronized seedling emergences are occasionally sown in seedbeds having unfavorable moisture due to lack of rainfall during planting period(Agigadi and Entz, 2002). For several years, strategies for enhancing the growth and development of crop species have been investigated for positive crop establishment, fast germination and emergence are essential, for which seed priming could play a pivotal role. Seed priming is a peculiar technology to promote rapid and uniform emergence and to achieve high vigor, resulting to better stand establishment and yield. Seed priming is a simple and low-cost hydration technique in which seeds are partially hydrated to an extent where pre-germination metabolic processestakeplace without actual germination and the seed is re-dried until close to the original dry weight. For better crop stand and high yields in a range of crop seedling priming is utilized.

Several studies have been carried out to reduce the period between sowing to emergence as this play a major role in crop production. Seed priming approach is one of the important developments in this area. The theory of seed priming was first proposed by Heydecker (1973). Harris *et al. (*2007) reported that seed priming yielded better establishment and growth, earlier flowering, higher seed tolerance to severe environmentalconditions and greater yield in maize. The merit of seed priming has been demonstrated for many field crops such as mung beans, barley, wheat, sweet corn lentil etc. (Sadeghian and yavari,2004).

Seed germination has been a triggeredconcern for several areasof theglobe; this is as a result of certain factors in the soil environment. The distribution of ferruginous soil all around the globe with high ironconcentrations which are more than the needed amount for plants to survive have been reported in severalrecent studies,a soil profile that is principally red and patchy yellow-red and possessstrong acid reactions.

Seed priming involve the technique of pre-soaking of seeds in a solution mainlyaid to breakdown seed dormancy and thereby facilitate germination capacity. The seeds of sorghum bicolor will be used for priming purpose. The novelty of this study is that it will enable us to know if seed priming technique promote germination of sorghum bicolor in a ferruginous soil. The aim of the study therefore is to enhance growth and germinability of sorghum under iron toxicity associated with ferrugenicity.

## Materials and methods

### Collection and preparation of material for the experiment

Seeds of sorghum (*Sorghum bicolor*) were obtained from the seed collection unit of the Office of the Agricultural Development Programme, Benin City, Edo State, Nigeria. Ferruginous (or iron elevated) soil used in this present study was obtained from around the Life Sciences Faculty environment and pooled to obtain composite sample. In order to confirm ferrugenicity, samples were collected from random areas and iron content was first confirmed in the area before more samples were collected and pooled. Further nutrient analyses were conducted on the sample and confirmed before soil was put to use (**Table 1**).

**Table 1:**
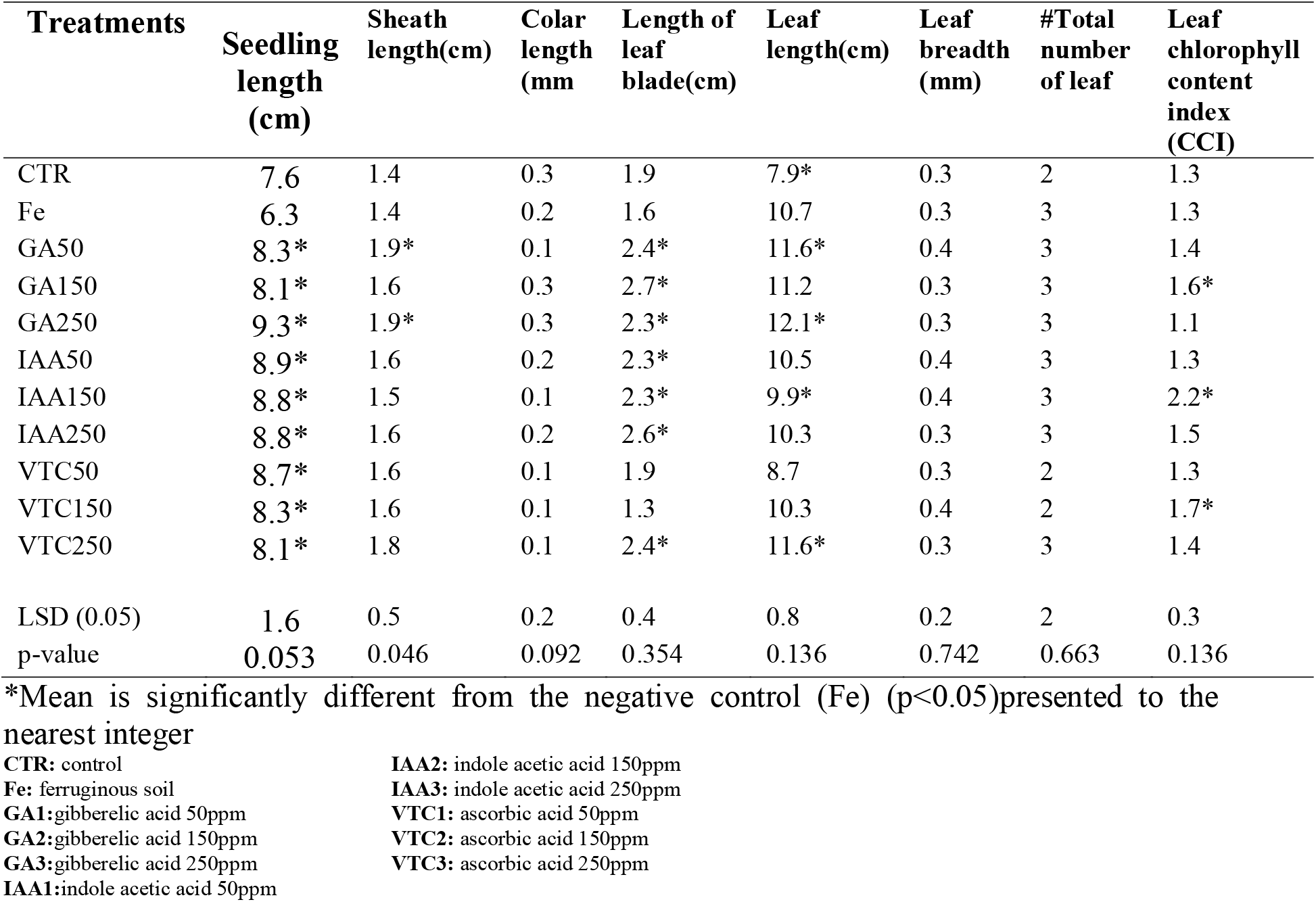
Shoot Growth Parameters at 7 days after Initiation.

### Soil Physiochemical Parameter

pH, Electric conductivity, Total organic carbon, Total Nitrogen, Exchangeable acidity, Clay, Silt, Sand and Iron (Fe) were determined in the laboratory according to the methods of APHA (2008).

### Soil Priming Material

The priming chemical agents used for the experiment were gibberellic acid, ascorbic acid and indole-acetic acid obtained in three (3) respective concentrations each (50, 150 and 250 ppm).

### Treatment and Experimental Design

Three priming media treatments were used for this experiment (GA: 50ppm, 150ppm, 250ppm; ascorbic: 50ppm, 150ppm, 250ppm;IAA: 50ppm, 150ppm, 250ppm). Water and ferruginous filtrate were used as control and negative control respectively. The duration of priming was set for three hours. The experiment was laid out incompletely randomized design in a factorial arrangement and replicated three times per treatment.

### Experimental Procedures

Top ferruginous soil was collected randomly from the marked plot, the soil was sieved to remove debris material, then water and soil was measured in ratio 1:1 by volume and stirred for about five minutes to havean evenly mixture. The mixture was decanted to separate filtrate from the sand. The sorghum seeds were immerged with solutions and primed for about three hours. Two layers of filter paper were placed in a 12cm petri dish. The filter paper was moistened with about 7ml of ferruginous filtrate, the sorghum seed is gently removed from their various solution and air dried. Twenty seeds were placed in two layers of filter paper in a 12cm petri dish and were left in the laboratory for about a week.

### Evaluations

#### Parameters Considered

The experimental set up, consisting of ferruginous soils in vitro and the priming seed were left in the laboratory for about one week, during this period; the following morphological parameters were assessed.

1. Number of germinated seeds: determined by physical counting.
2. Shoot length: determined by using a metric rule.
3. Root length(cm): determined by using a meter rule.
4. Sheath length(cm): determined by using a meter rule.
5. Colar length(cm): determined by using a meter rule.
6. Length of leaf blade(cm): determined by using a meter rule.
7. Leaf length(cm): determined by using a meter rule.
8. Leaf breadth(mm): determined by using a meter rule.
9. Number of roots: determined by physical counting.
10. Total number of leaves: determined by physical counting.
11. Number of nodes: determined by physical counting.
12. Chlorophyll content of the leaf(ccl): determined by using chlorophyll meter.
13. Plant wet weight(g): determined by using electric weight meter.
14. Plant dry weight(g): determined by using electric weight meter.

### Root and Shoot growth

One seedling waschosen among the prominent height randomly from petri dish after germination test. Root and shoot were excised from the seedling and shoot and root length was measured with a transparent meter rule. The seedling was weighed to determine fresh weight. The seedling dry weight was measured after drying ofthe samples exposed to air.

### Assessment of Germination Indices

#### First day of germination (or Germinability) (FDG)

It is the time when the first germination was recorded.

#### Last day of germination (LDG)

This is the last day when the seed germination was reported.

#### Finalgermination percentage (FGP)

This is the germination percentage attained by the plant even beyond the time period.

#### Peak period of germination (PPG) or Modal time of germination (MTG)

It is the time in which highest frequency of germinated seeds are observed and

#### Median germination time (MeGT), or Days required for 50% germination, T_50_

Days required for 50% germination, T_50_

T_50_ = Days required for 50% germination of the total number of seeds

#### Germination rate index (GRI)

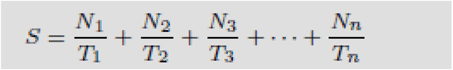

Where GP_1_ is germination percentage at 1^st^ day, GP_2_ is germination percentage at 2 days,GP_n_ is germination percentage at n days. GRI reflects the percentage of germination on each day of the germination period.

#### Speed of accumulated germination, SAG

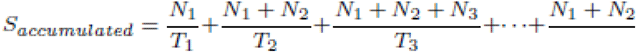

Where GP_1_ is germination percentage at 1^st^ day, GP_2_ is germination percentage at 2 days,GP_n_ is germination percentage at n days. GRI reflects the percentage of germination on each day of the germination period.

#### Corrected germination rate index (GRI_corrected_)

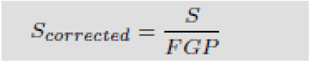

#### Timson’s Index, (TI) or Germination Energy Index

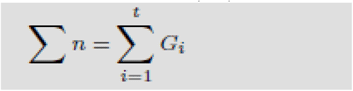

Where GP1, GP2, …GPn are the germination percentages at day 1, 2, … and *n* respectively.

#### Modified Timson’s Index, (TI_mod_)

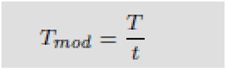

#### The seedling vigor

SVI (I) = Seedling length x FGP

SVI (II) = (Root length + Shoot length) x FGP

SVI (III) = Seedling dry weight x FGP

#### Time spread of germination, or Germination distribution (TSG)

TSG = LDG - FDG

This is the time (in days) taken between the first and last germination events.

#### Germination index, GI

**GI = 10 x (S**_**1**_**+S**_**2**_**+S**_**3**_**+ … +S**_**n**_**) / (1*S**_**1**_ **+ 2*S**_**2**_ **+ 3*S**_**3**_ **+ … + *n* * S**_***n***_)

WhereS_1_, S_2_, S_3_,Sn are number of seeds that germinated per lot (or petri dish) at day 1, day 2, day 3 … day n

#### Mean dailygermination, MDG

MDG = FGP/d, where d is the number of days it took to first arrive at the FGP

#### Daily germination speed, DGS

DGS = 1 /MDG. This is the reciprocal of MDG

#### Mean germination time, MGT

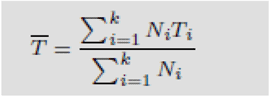

Where S_1_, S_2_, S_3_,Sn are number of seeds that germinated per lot (or petri dish) at day 1, day 2, day 3, …day n.The lower the MGT, the faster a seed population has germinated.It is also called Length of Germination Time (LGT) or Germination Resistance (GR) or Sprouting Index (SI). It is the average length of time required for maximum germination of a seed lot.

#### Germination Value (Czabator)

GV = PV x MDG

Where, PV is the peak value and MDG is the mean daily germination percentage from the onset of germination.

#### Mean germination rate, MGR

MGR = 1 / MGT

#### Coefficientof velocity ofgermination, CVG

CVG = [(G_1_+G_2_+G_3_+ … +G_n_) / (1*G_1_ + 2*G_2_ + 3*G_3_ + … + *n* * G_*n*_)] * 100

Where G_1_, G_2_, G_3_,Gn are germination percent per lot (or petri dish) at day 1, day 2, day 3 … day n. CVG gives an indication of the rapidity of germination.

Or, simply, CVG = MGR x 100

#### Variance of germination time, VGT or S^2^_T_

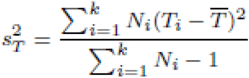

Where, N_1_, N_2_, N_3_, … and N*i* are the number of seeds germinated in 1st, 2nd, 3rd, … and *i*th day respectively

#### Germination capacity, GC

GC = FGP / N

Where N is number of seeds used in the bioassay

#### Variance of germination rate, VGR

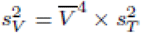

Where, 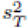 is the variance of germination time.

VGR = (MGR)^4^ x VGT

#### Synchronization Index

Or Uncertainty of the germination process

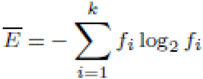

Where, *f*_*i*_ is the relative frequency of germination 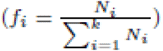, *N*_*i*_ is the number of seeds germinated on the *i*th time and *k* is the last day of observation.

### Data analysis

The complete randomized design was done using the Microsoft excel-2010 statistical software. The mean significant difference was separated from the least significant difference.

**Plate 1:**
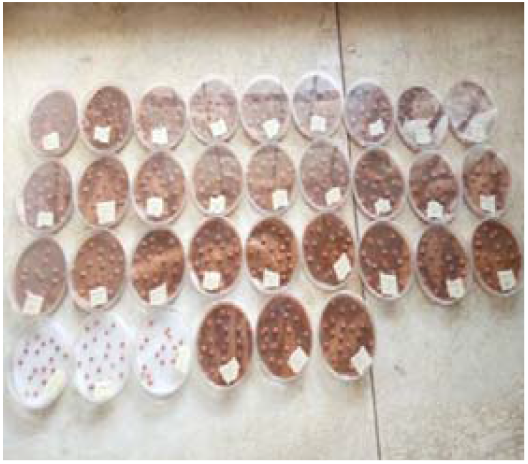
Seed emerged in ferruginous solution.

**Plate 2:**
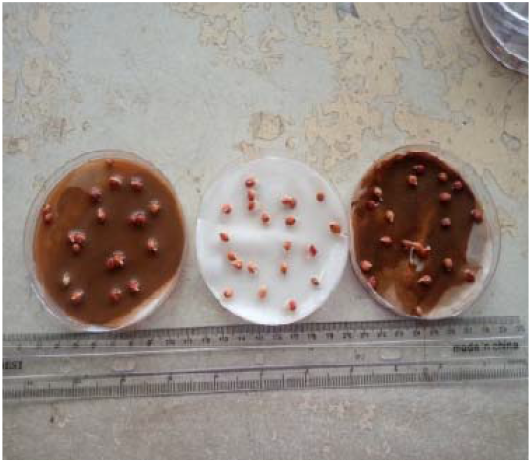
Plumule emerged at day 2.

**Plate 3:**
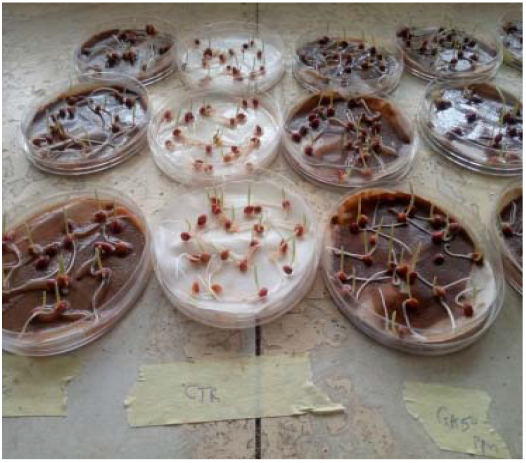
Primary root seen at day 3

## Results

Table 1 shows the soil characteristics of soil used in the study. The soil was slightly acidic (pH = 5.39) and was somewhat reddish brown in colour. The iron content of soil was 382.09 mg/kg for which it was termed ferruginous.

**Table 1:** Physical and Chemical Properties of soil used in the Study.

| Parameters | Soil (n = 3) |
| --- | --- |
| pH | 5.39 |
| Electrical conductivity (μs/cm) | 311.72 |
| Total organic carbon (%) | 0.51 |
| Total Nitrogen (%) | 0.19 |
| Exchangeable acidity (meq/100 g soil) | 0.22 |
| Na (meq/100 g soil) | 9.93 |
| K (meq/100 g soil) | 1.22 |
| Ca (meq/100 g soil) | 16.01 |
| Mg (meq/100 g soil) | 12.75 |
| Clay (%) | 5.24 |
| Silt (%) | 9.06 |
| Sand (%) | 85.70 |
| Fe (mg/kg) | 382.09 |

Figure1 shows the germination percentage of the test plant in gibberellic acid, indole-acetic acid and ascorbic acid. Result shown that germination percentage ranged from 75% at first day after germination initiation to 95% on the fourth day after germination when the seeds were sub-emerged in gibberellic acid. Generally, germination percentage in ferruginous medium was significantly reduced; in the first day germination percentage was 75% and 90% on the third day after germination initiation. When seeds were exposed to indole acetic acid germination percentage ranged from 80% to 95% in four days compared to 75% to 95% in the same four days when seed were exposed to ascorbic acid. Generally, the highlight of the application of the growth stimulator was lag although they was significant reduction in germination percentage when seeds were exposed to ferruginous solution compared to the control; however, the application of gibberellic acid mostly enhanced germination percentage then indole acetic acid and ascorbic acid; germination percentage after fourth day was 80% and at the fifth day it was already 100% for at least gibberellic acid (GA50). However, the highlight for indole acetic acid was that it most likely reduced germination time compared to gibberellic acid and ascorbic acid; time taken to achieved 100% germination percentage was just 48 hours at least in indole acetic acid (IAA50) (**Figure 1. b**).

**Figure 1:**
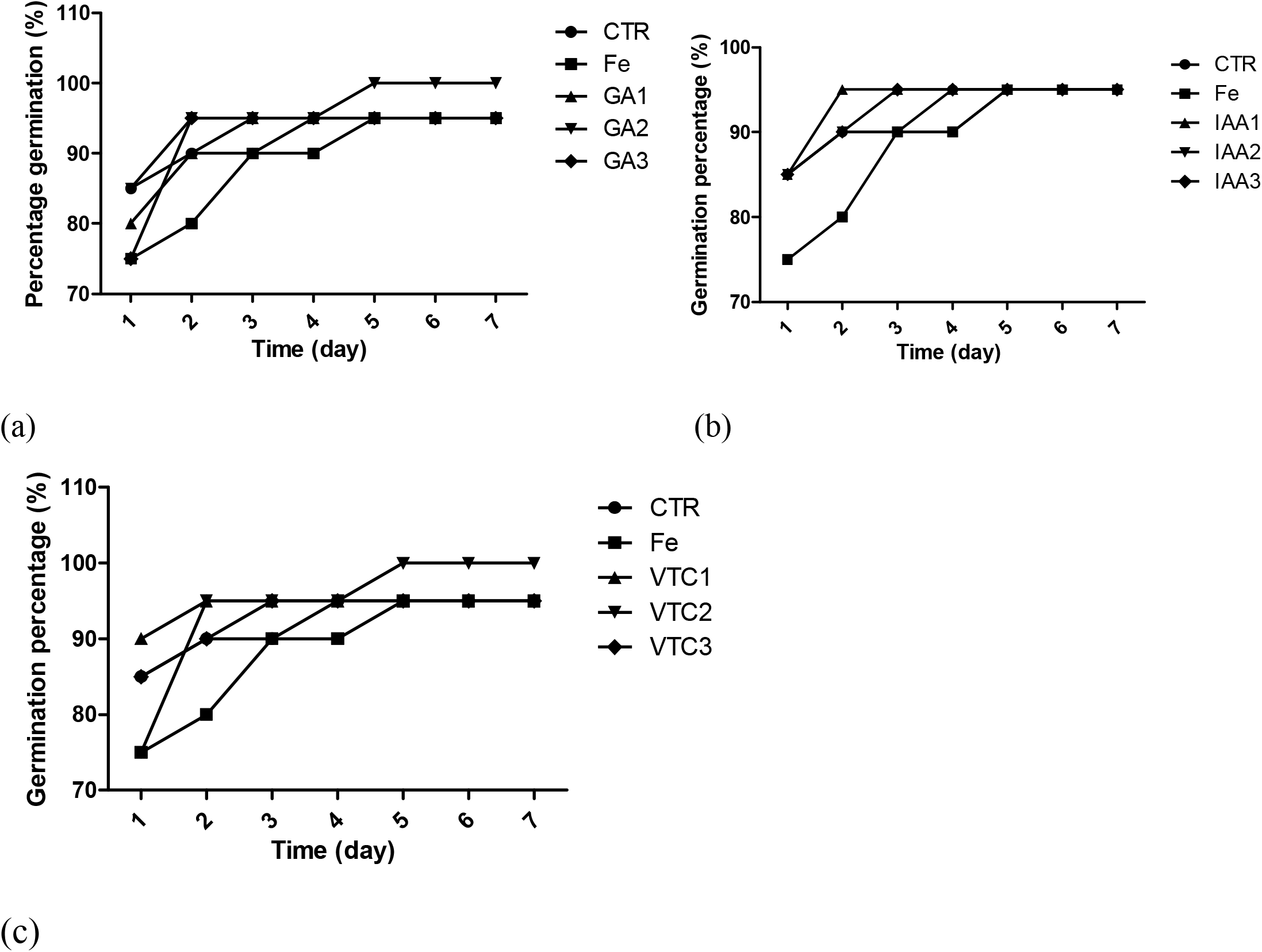
Germination percentage of test plant in (a) gibberellic acid (b) indole acetic acid and (c) ascorbic acid exposed seeds in a ferruginous medium **Key:** CTR: control Fe: ferruginous soil GA1: gibberelic acid 50ppm GA2: gibberelic acid 150ppm GA3:gibberelic acid 250ppm IAA1:indole acetic acid 50ppm IAA2: indole acetic acid 150ppm IAA3: indole acetic acid 250ppm VTC1: ascorbic acid 50ppm VTC2: ascorbic acid 150ppm VTC3: ascorbic acid 250ppm

**Figure 2:**
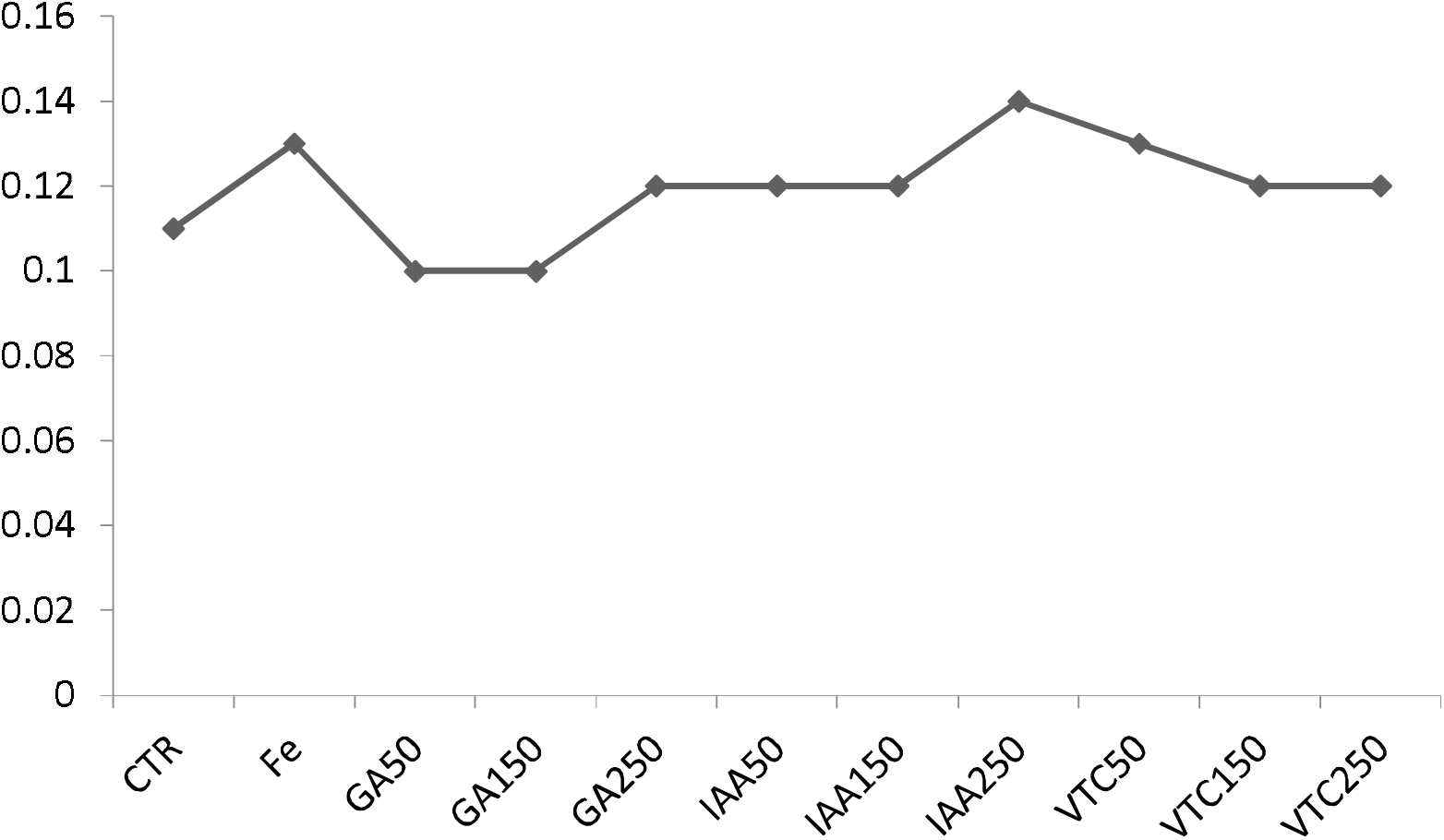
Five (5) seedling dry weight of Sorghumat 7 days after germination initiation

There was no significant difference in the 5 seedlings weight of Sorghum, as values ranged between 0.11 to 0.14 grams (**Figure 2**).

The shoot growth parameter of Sorghum at 7 days after germination initiation was presented on Table 1. Result shown that, although they was reduction in seedling length in the seed exposed to ferruginous solution without the application of growth stimulators (6.3cm), it was not significantly different from that of the control (7.6cm) (P>0.05). However, when the growth stimulators gibberellic acid, indole-acetic acid and ascorbic acid were applied at various concentration, 50, 150, and 250 ppm, seedling height was significantly enhanced compared to the values obtained in the control and value obtained from ferruginous solution (8.1 to 9.3cm) (P,0.05), similarly sheath length in gibberellic acid enhanced plant in ferruginous solution was significantly higher (1.9cm) when compared with the negative control (1.4cm). In this study whereas the control solution didnot receive ferruginous application, the negative control had those seeds exposed to ferruginous solution but were not administered the three growth stimulating agents. Generally, they was no significant difference in colar length (0.1 to 0.3mm), leaf breadth (0.3 to 0.4mm) and the total number of leaf after 7 days (2 to 3cm). However, the higher concentration of growth stimulator significantly promoted leaf chlorophyll content index when compared to both control and negative control (**Table 2**).

**Table 2:** Root Growth Parameters at 7 days after Initiation.

| Treatments | Root length (cm) | Total number of root branching |
| --- | --- | --- |
| CTR | 7.9* | 5 |
| Fe | 9.7 | 6 |
| GA1 | 7.3* | 7 |
| GA2 | 7.3* | 8 |
| GA3 | 7.4* | 6 |
| IAA1 | 8.7 | 5 |
| IAA2 | 8.9 | 9 |
| IAA3 | 5.7* | 10* |
| VTC1 | 6.1* | 9 |
| VTC2 | 5.1* | 9 |
| VTC3 | 6.3 | 8 |
| LSD (0.05) | 1.4 | 4 |
| p-value | 0.029 | 0.009 |

\*Mean is significantly different from the negative control (Fe) ( $p < 0.05$ )
CTR: control
Fe: ferruginous soil
GA1: gibberelic acid 50ppm
GA2: gibberelic acid 150ppm
GA3: gibberelic acid 250ppm
IAA1: indole acetic acid 50ppm
IAA2: indole acetic acid 150ppm
IAA3: indole acetic acid 250ppm
VTC1: ascorbic acid 50ppm

When considering the roots parameter, it was observed that shoot length were significantly reduced in plant exposed to ferruginous solution without growth stimulating agentsbut the root parameter were enhanced (**Table 2**). Root length in ferruginous solution was (9.7cm), however, when the seeds were exposed to the three-growth stimulating agents there was significant reduction in root length from (5.1 to 8.9cm, P<0.05). However, they were no significant changes in the total number of root branches irrespective of ferrugenic status or application of growth stimulating agents (**Table 2**).

Presentation of germination time in germination indices is shown in (Table 3). The result showed that it took between 19.6 hours to 32.1 hours for first day of germination to occur in Sorghum. Result also showed that when seeds were subjected to ferrugenicity, germinability was 31.2 hours compared to 22.3 hours in the control.However, when the seeds in ferruginous solution were enhanced with growth stimulators, they was significant reduction in germinability from 31.2 hours to between 19.6 hours and 21.1 hours in GA50,GA250, GA150, IAA150, VTC50, and VTC150 ppm respectively.

**Table 3:** Measurement of Germination Time.

| Parameters | CTR | Fe | GA1 | GA2 | GA3 | IAA1 | IAA2 | IAA3 | VTC1 | VTC2 | VTC3 | LSD(0.05) | p-value |
| --- | --- | --- | --- | --- | --- | --- | --- | --- | --- | --- | --- | --- | --- |
| First day of germination (or Germinability) (FDG) (hr) | 22.3* | 31.2 | 22.4* | 23.1 | 21.2* | 32.1 | 22.4* | 25.4 | 22.1* | 19.6* | 31.2 | 8.4 | 0.302 |
| Last time of germination (LDG) (hr) | 71.9 | 71.9 | 61.2 | 61.2 | 63.9 | 50.2* | 35.1* | 34.5* | 71.9 | 49.0* | 63.9 | 12.4 | 0.004 |
| Time spread of germination, or Germination distribution (TSG) (hr) | 51.3 | 43.2 | 39.4 | 42.3 | 45.4 | 24.2* | 12.3* | 13.2* | 51.3 | 31.2 | 34 | 14.7 | 0.035 |
| Peak period of germination (PPG) or Modal time of germination (MTG) | 71.9 | 71.9 | 71.9 | 52.3* | 42.3* | 51.3* | 34.0* | 35.4* | 71.9 | 49.0* | 36.4* | 18.4 | 0.028 |
| Median germination time (MeGT), or Days required for 50% germination, T<sub>50</sub> | 22.5 | 24.3 | 21.5* | 24.3 | 22.5 | 26.4 | 21.5* | 24.6 | 25.3 | 23 | 25 | 2.5 | 0.135 |
| Germination rate index (GRI) | 183.8 | 186.5 | 156.7 | 207.9 | 122.5 | 132.5 | 201.5 | 130.0 | 137.5 | 197.9 | 161.7 | NA | NA |
| Mean daily germination, MDG | 23.8 | 19.0 | 31.7 | 20.0 | 47.5 | 47.5 | 19.0 | 31.7 | 47.5 | 20.0 | 31.7 | NA | NA |
| Daily germination speed, DGS | 0.0 | 0.1 | 0.0 | 0.1 | 0.0 | 0.0 | 0.1 | 0.0 | 0.0 | 0.1 | 0.0 | NA | NA |
| Mean germination time, MGT | 2.5 | 3.1 | 2.1 | 3.1 | 1.6 | 1.5 | 3.0 | 2.0 | 1.5 | 3.1 | 2.0 | NA | NA |
| Mean germination rate, MGR | 0.4 | 0.3 | 0.5 | 0.3 | 0.6 | 0.7 | 0.3 | 0.5 | 0.7 | 0.3 | 0.5 | NA | NA |
| Speed of accumulated germination, SAG | 350.8 | 403.9 | 253.3 | 453.2 | 160.0 | 175.0 | 439.6 | 262.5 | 182.5 | 430.3 | 262.5 | NA | NA |
| Corrected germination rate index (GRI<sub>corrected</sub>) | 1.9 | 2.0 | 1.6 | 2.1 | 1.3 | 1.4 | 2.1 | 1.4 | 1.4 | 2.0 | 1.7 | NA | NA |
| Coefficient of velocity of germination, CVG | 39.3 | 32.1 | 48.6 | 32.6 | 64.2 | 65.5 | 32.8 | 49.1 | 66.1 | 32.2 | 49.1 | NA | NA |

\*Statistically different from the negative control (Fe)
**CTR:** control
**Fe:** ferruginous soil
**GA1:** gibberelic acid 50ppm
**GA2:** gibberelic acid 150ppm
**GA3:** gibberelic acid 250ppm
**IAA1:** indole acetic acid 50ppm
**IAA2:** indole acetic acid 150ppm
**IAA3:** indole acetic acid 250ppm
**VTC1:** ascorbic acid 50ppm

The last time of germination in seeds exposure to ferrugenicity was 71.9 hours. However, when these seeds were exposed to indole acetic acid and ascorbic acid, last time of germination was reduced significantly from 71.9 hours to between 34.5 hours and 49.0 hours IAA50, IAA150, IAA250, and VTC150 respectively. That ordinarily implied that germination distribution in time was significantly reduced by the application of indole acetic acid from 43 hours in seeds exposed to ferrugenicity to between 12.3 and 24.2 hours (P<0.05). Generally, also the peak period of germination or modal time of germination which indicated the time when the higher germination percentage was first recorded in the experiment occurred with seeds treated with GA150 (52.3 hours), GA250 (43.3 hours), IAA50 (51.3 hours), IAA150 (34.0 hours), IAA250 (34.4 hours), VTC150 (49.0 hours), and VTC250 (36.4 hours) respectively. Germination rate index were lower in the seeds that were exposed to ferrugenicity but latter enhanced with growth stimulator when compared with those that ordinarily left in ferruginous solution.

The measurements of germination capacity were achieved in Table 4. Timson’s index which measured germination energy was high in seed exposed to ferrugenicity (1340). However, with the application of growth stimulators, germination energy reduced to as low as (265) in gibberellic acid 250 ppm. Result from modifiedTimson’s index showed that seedling in ferruginous solution was 268 compared to 181.7 index when seeds were exposed to gibberellic acid 50ppm and 132.5 when seeds were exposed to gibberellic acid 250 ppm. Germination value were significantly low in seeds exposed to ferruginous (1805.0) compared to the control (2256.3). However, application of the growth stimulator up to at least 100ppm increase germination value to values above 2000. However, no difference observed in the germination capacity as value ranged between 4.8 to 5.0 (**Table 4**).

**Table 4:** Measurement of Germination Capacity.

| Parameters | CTR | Fe | GA1 | GA2 | GA3 | IAA1 | IAA2 | IAA3 | VTC1 | VTC2 | VTC3 |
| --- | --- | --- | --- | --- | --- | --- | --- | --- | --- | --- | --- |
| Timson's Index, (TI) or Germination Energy Index | 915.0 | 1340.0 | 545.0 | 48060.0 | 265.0 | 275.0 | 1370.0 | 550.0 | 280.0 | 1430.0 | 550.0 |
| Modified Timson's Index, (TI<sub>mod</sub>) | 228.8 | 268.0 | 181.7 | 9612.0 | 132.5 | 137.5 | 274.0 | 183.3 | 140.0 | 286.0 | 183.3 |
| Germination Value (Czabator) | 2256.3 | 1805.0 | 3008.3 | 2000.0 | 4512.5 | 4512.5 | 1805.0 | 3008.3 | 4512.5 | 2000.0 | 3008.3 |
| Germination capacity, GC | 4.8 | 4.8 | 4.8 | 5.0 | 4.8 | 4.8 | 4.8 | 4.8 | 4.8 | 5.0 | 4.8 |

Table 5 showed the measurements of seedling vigor due to treatment exposure, result showed very low seedling vigor using seedling vigor 1 whichdetermined by (Baki and Andersen, 1972). Reduced seedling vigor in the seed exposed to ferrugenicity (598.5)were eventually enhanced when these same seeds were treated with either gibberellic acid, indole-acetic acid and ascorbic acid during where seedling vigor were increased by more than 50% compared to initial value in ferrugenicity (Table 5). However, minimal difference in seedling vigor was obtained when vigor was determined as a factor of seedling weight. Since earlier result showed no significant difference in seedling weight, seedling vigor III as presented on table 5 indicated minimal difference as value ranged from 2.9 in IAA50 to 5.0 in both GA150 and VTC 150; whereas values in ferruginous solution without stimulator was 3.8.

**Table 5:** Measurement of Seedling Vigour due to Treatment Exposure.

| Parameters | CTR | Fe | GA1 | GA2 | GA3 | IAA1 | IAA2 | IAA3 | VTC1 | VTC2 | VTC3 |
| --- | --- | --- | --- | --- | --- | --- | --- | --- | --- | --- | --- |
| The seedling vigor, I | 703.0 | 598.5 | 788.5 | 810.0 | 883.5 | 807.5 | 836.0 | 836.0 | 703.0 | 830.0 | 760.0 |
| The seedling vigor, II | 1168.5 | 1349.0 | 1235.0 | 1240.0 | 1301.5 | 1349.0 | 1396.5 | 1092.5 | 997.5 | 1040.0 | 1111.5 |
| The seedling vigor, III | 3.8 | 3.8 | 4.8 | 5.0 | 4.8 | 2.9 | 2.9 | 3.8 | 4.8 | 5.0 | 3.8 |

The synchronization index in any germination experiment is very important because it show the possibility that germination will not occur. When synchronization index is low that implies the higher capacity indicative that germination may not occur. In table 6, synchronization index presented revealed an index of 73.2 when seedswere exposed to ferrugenicity compared to 87.5 in control. This ordinarily implies that the presence of ferruginous solution hasasignificantly impeding impact on the germination performance of Sorghum. However, in spite of the impeding capacity of ferruginous nature of the solution used in this germination experiment, synchronization was significantly increased from 73.20 to as high as 89.92 when there were exposed growth stimulating agents. This implies therefore that in spite of the impudence attributed to ferrugenicity, the application of growth stimulators enhanced the capacity of germination. Plates 4 and 5 show seedlings on day 6.

**Table 6:** Synchronization Index.

| Time (days) | Ni | Fi | $\log_2(\text{Fi})$ | $-(\text{Fi}) * \log_2(\text{Fi})$ |
| --- | --- | --- | --- | --- |
| **Control** |  |  |  |  |
| 1 | 17 | 17.00 | 4.09 | 68.49 |
| 2 | 18 | 6.00 | 2.58 | 14.51 |
| 3 | 18 | 3.00 | 1.58 | 3.75 |
| 4 | 19 | 1.90 | 0.93 | 0.76 |
|  |  |  | *Total* | 87.51 |
| **Fe** |  |  |  |  |
| 1 | 15 | 15.00 | 3.91 | 57.60 |
| 2 | 16 | 5.33 | 2.42 | 11.88 |
| 3 | 18 | 3.00 | 1.58 | 3.75 |
| 4 | 18 | 1.80 | 0.85 | 0.53 |
| 5 | 19 | 1.27 | 0.34 | -0.57 |
|  |  |  | *Total* | 73.20 |
| **GA50** |  |  |  |  |
| 1 | 16 | 16.00 | 4.00 | 63.00 |
| 2 | 18 | 6.00 | 2.58 | 14.51 |
| 3 | 19 | 3.17 | 1.66 | 4.27 |
|  |  |  | *Total* | 81.78 |
| **GA150** |  |  |  |  |
| 1 | 17 | 17.00 | 4.09 | 68.49 |
| 2 | 19 | 6.33 | 2.66 | 15.87 |
| 3 | 19 | 3.17 | 1.66 | 4.27 |
| 4 | 19 | 1.90 | 0.93 | 0.76 |
| 5 | 20 | 1.33 | 0.42 | -0.45 |
|  |  |  | *Total* | 88.93 |
| **GA250** |  |  |  |  |
| 1 | 15 | 15.00 | 3.91 | 57.60 |
| 2 | 19 | 6.33 | 2.66 | 15.87 |
|  |  |  | *Total* | 73.47 |
| **IAA50** |  |  |  |  |
| 1 | 17 | 17.00 | 4.09 | 68.49 |
| 2 | 19 | 6.33 | 2.66 | 15.87 |
|  |  |  | *Total* | 84.35 |
| **IAA150** |  |  |  |  |
| 1 | 17 | 17.00 | 4.09 | 68.49 |
| 2 | 18 | 6.00 | 2.58 | 14.51 |
| 3 | 18 | 3.00 | 1.58 | 3.75 |
| 4 | 18 | 1.80 | 0.85 | 0.53 |
| 5 | 19 | 1.27 | 0.34 | -0.57 |
|  |  |  | *Total* | 86.71 |
| **IAA250** |  |  |  |  |
| 1 | 17 | 17.00 | 4.09 | 68.49 |
| 2 | 18 | 6.00 | 2.58 | 14.51 |
| 3 | 19 | 3.17 | 1.66 | 4.27 |
|  |  |  | *Total* | 87.26 |
| **VTC50** |  |  |  |  |
| 1 | 18 | 18.00 | 4.17 | 74.06 |
| 2 | 19 | 6.33 | 2.66 | 15.87 |
|  |  |  | *Total* | 89.92 |
| **VTC150** |  |  |  |  |
| 1 | 15 | 15.00 | 3.91 | 57.60 |
| 2 | 19 | 6.33 | 2.66 | 15.87 |
| 3 | 19 | 3.17 | 1.66 | 4.27 |
| 4 | 19 | 1.90 | 0.93 | 0.76 |
| 5 | 20 | 1.33 | 0.42 | -0.45 |
|  |  |  | *Total* | 78.05 |
| **VTC250** |  |  |  |  |
| 1 | 17 | 17.00 | 4.09 | 68.49 |
| 2 | 18 | 6.00 | 2.58 | 14.51 |
| 3 | 19 | 3.17 | 1.66 | 4.27 |
|  |  |  | *Total* | 87.26 |

**Plate 4:**
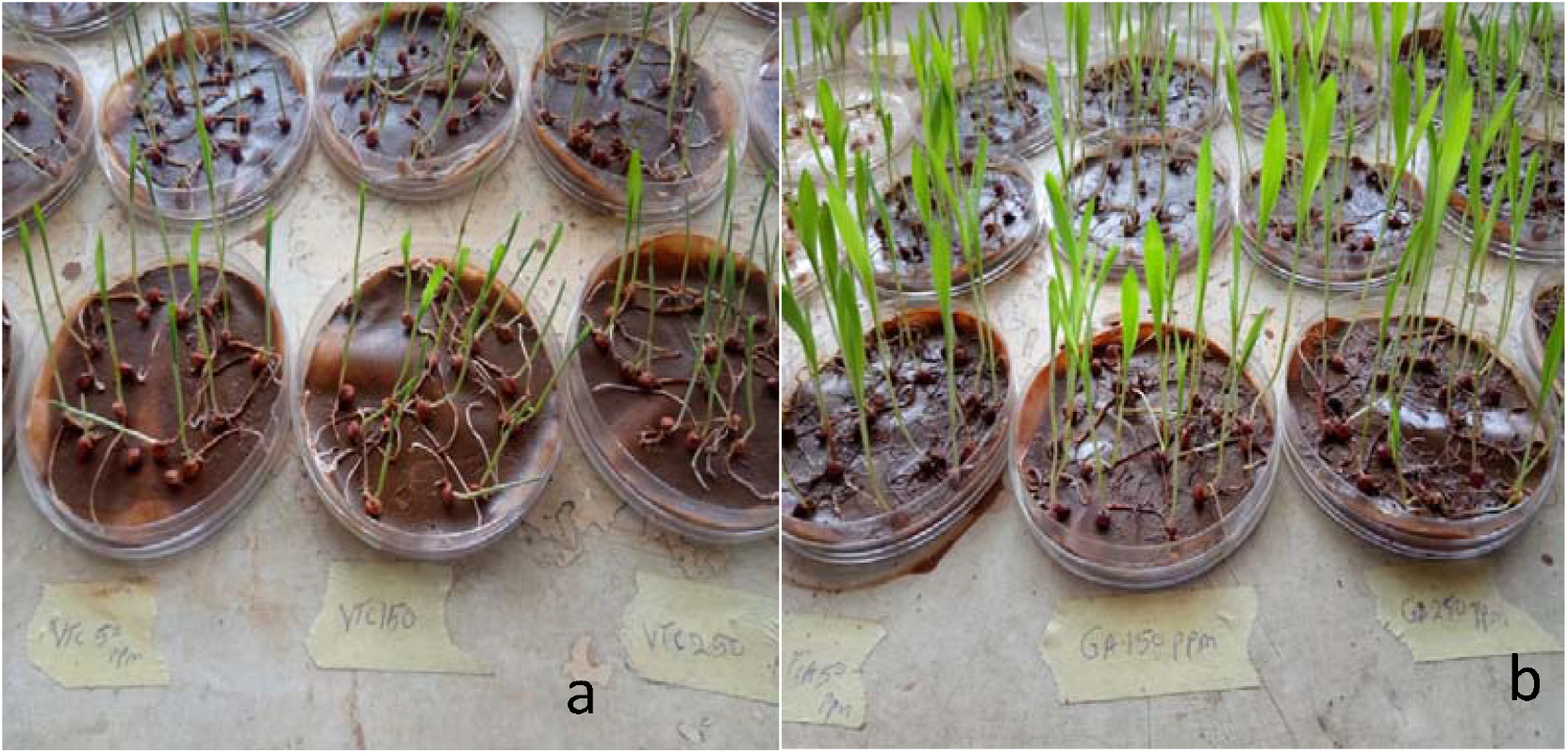
(a)Seedling at day 4. (b) Seedling at day 5

**Plate 5:**
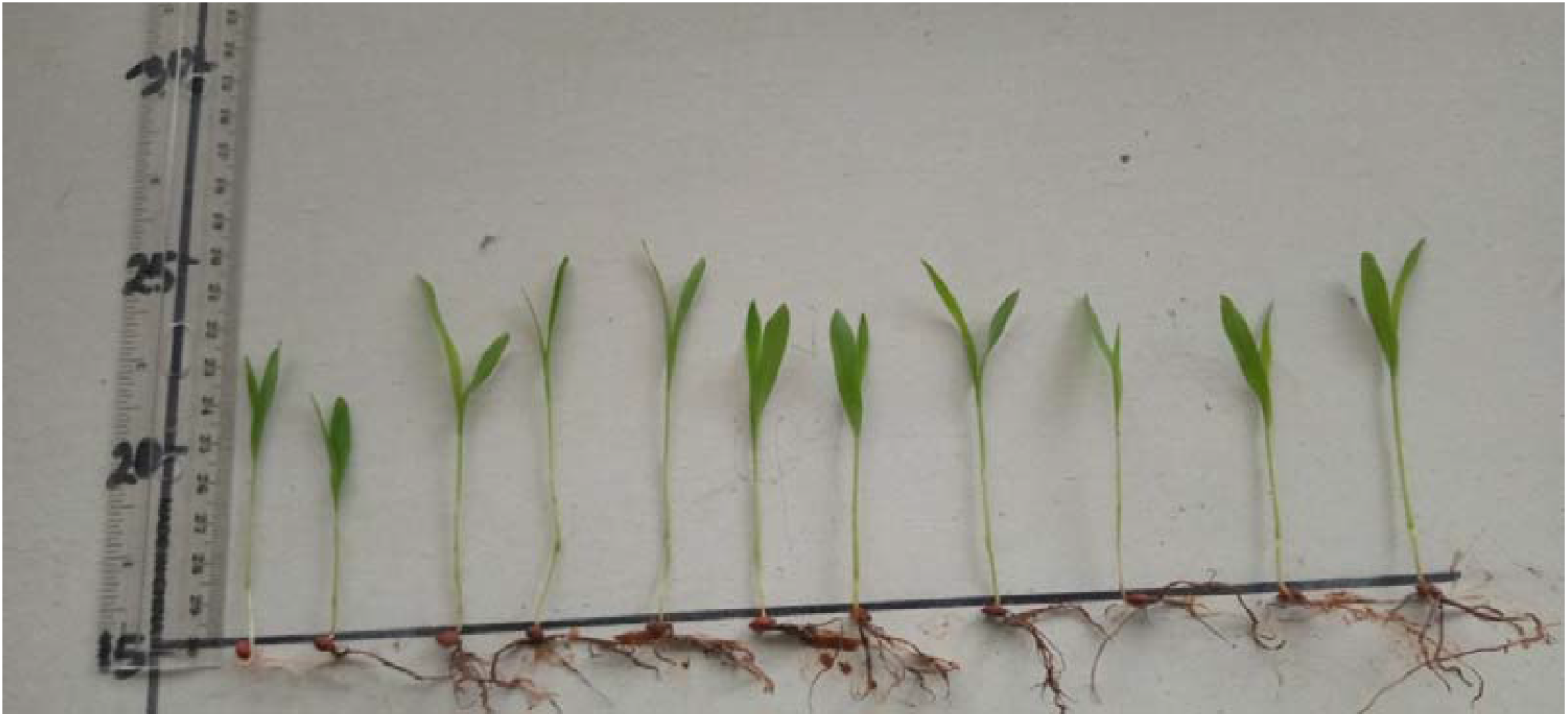
Sorghum plant placed at whiteboard to determine the height of each treatment **Key**: left to right Control(water), ferruginous, gibberellic acid 50ppm, gibberellic acid 150ppm, gibberellic acid 250ppm, indole acetic 50ppm, indole acetic 150ppm, indole acetic 250ppm, ascorbic 50ppm, ascorbic 150ppm, ascorbic 250ppm.

## Discussion

The presence of iron toxicity in plants is linked with increasedFe^2+^ uptake by roots and its movement to leaves through transpiration stream. ExcessiveFe^2+^ causes free radical production that hampers cellular structure irredeemably and destroys membranes, DNA and proteins (Arora et al., 2002; Dorlodot *et al*., 2005). Iron toxicity in tobacco, canola, Soybean and *Hydrillaverticillata* are followed with adecrease of plant photosynthesis, yield and increase of oxidative stress and ascorbate peroxidase activity (Kampfenkel and Montagu, 1995; Caro and Puntarulo, 1996).

There was significant reduction in germination indices with ferrugenicity. In seedling exposed to ferruginous solution the pace of root growth (measured in cm) was increased considerably in ferruginous treatment after germination initiation. It was obvious from the data that there was a threshold Fe concentration above which the growth of Sorghum seedlings was seriously inhibited.Root growth measurement in cm confirmed the inhibitory impact of elevated concentrations of Fe.

Seedlings exposed to ferrugenicityreally showed stimulation of root growth. These seedlings suffered a decline in Fe supply; hence the plants may have reallocated resources to increase root growth in a bidto locate new sources of this vital nutrient. Previous researchstudiesof wetland species including *P. australis*, have shown glaring effects of high iron concentrations liken to those found here (Snowden and Wheeler, 1995). Generally red soils are usually poor growing soil with low plant yield (Yu et al., 2016).

Even though available iron in solution reduced germinability of seeds, introduced growth stimulator facilitated the germination outcome. This is even more obvious in the synchronization index. Poor and unsynchronized seedling emergences are occasionally sown in seedbeds having unfavourable moisture as a result of lack of rainfall at sowing time (Agigadi and Entz, 2002). Plant growth stimulators in this present study had shown to enhance seedling index in ferruginoussoilin vitro, as similar study showed that GA3 and Ethrelenhanced the seed quality and showed improved seedling length, seedling dry weight which in turn improved higher seedling vigour index, germination speed and mean germination time (Sarika *et al*., 2013). Plant growth stimulators appear to play a paramount role in the source-sink relations in the plant. Gibberellic acid promotes the sink activity of the shoot, gibberellic acid is among the most widely utilized plant growth regulators which facilitates stem elongation along with plant height dry matter build-up, growth as well as yield in various crops (Harrington et al.,1996; Akter *et al*., 2007 and Emongor,2007). Hence, the present investigation was carried out to study the effect of gibberellic acid, indole acetic acid and ascorbic acid on plant growth, dry matter and yield parameter of Sorghum. Growth stimulator snowballed the growth activities of theplant, as the result showed that there was a compensatory growth between the shoot length and the root length (Table 1 and 2) as this is more in IAA. Similar studies had been done in *Hibiscus sabdariff (*Ali et al., (2012).

Overall indole acetic acid produced the best germination response in all concentrations utilized. An experiment carriedout in cowpea seeds showed that indole acetic acid (IAA) accelerated seed germination percentage reaching value of 85% and 89% respectively (Luma *et al*., 2019), which is more or likely the same when seeds were exposed to indole acetic acid germinationas percentage ranged from 80% to 95% in four days compared to 75% to 90% in the same four days when seed were exposed to ferruginous solution (Figure 1). A study in barley cultivars showed that indole acetic acid successfully promoted plant growth and reduced adverse effect of water stress (Ashraf et al., 2006) as it shown in this study with seeds exposed to ferruginous solution which happened to contain iron.

As an essential stimulator, IAA plays a vital role in the growth and development of cotton plants (Min et al., 2014; Zhang et al., 2011). In this study, it was shown that germination rate, index, and speed were all significantly enhanced after IAA priming in Sorghum. These results are in consonancewith the previous research that seed priming by chemicals can facilitate seed germination (Espanany *et al*., 2016;Rahimi, 2013; Seyyedi *et al*., 2015). The study also revealed that IAA stimulator had an advantageouseffect on both the initial germination and the continued growth of Sorghum seedlings suggesting that IAA used as seed stimulator may increase stand formation and eventually promote plant production. Compared with germination stage, biomass including fresh and dry weight was significantly more built-up in IAA than ferruginous solution in seedling stage, which might bebecause photosynthesis was more activated by IAA stimulator. During the germination stage, IAA priming might cause more fresh weight and size based on water uptake, but the dry weight was still the same. Previously, some studies were carried out to address the impact of seed priming on seedling growth, and concluded that different chemicals could increase seedling growth by different mechanism.

Priming with melatonin, the root and plant weight snowballed under salt stress in basil by counteracting the effects of salinity regarding metabolites and promoting antioxidant capacity (Bahcesular *et al*.,2020). In this study, it was portrayed that IAA priming could enhance the seedling growth in Sorghum by modulating the endogenous phytohormones synthesis and balance.

Priming with polyethylene glycol, silicon, and selenium could significantly enhance performance during water stresscondition (Nawaz et al., 2012; Paryeen *et al*., 2019; Rahimi, 2013). Priming with melatonin and CaCl_2_, the tolerance against salt stress were significantly enhanced (Bahcesular *et al*., 2020; Tabassum et al., 2017). Seed priming technique also could significantly enhance the germination rate and germination average time (Espanany *et al*., 2016; Nakaune *et al*., 2012; Rahimi 2013), which has a huge positive significance in agriculture production. The result in the present study showed that IAA priming could significantly enhance seed germination and seedling vigor, which could aid plants to meet the demands of undesirable growing conditions, such as cold and drought stress after seed sowing, as quicker seed germination and stronger seedling vigor would permit early planting and rapid establishment of seedlings to have a longer span of reproductive growth and potentially anincreased yield.

In plants, it has beenobserved that the most essential antioxidant is ascorbate which in combination with other defense system components, safe guidesplants against oxidative impairment resulting from aerobic metabolism, photosynthesis and avariety of pollutants.It inhibits membrane peroxidation, thereby shielding cell from harm and prolonging their scenescene (Zhang,2013).Ascorbic acid affects several physiological mechanisms, including plant metabolic differentiation and increased availability of water and nutrients, thus enabling plants to be guided against environmental stresses such as salt stress (Khan et al., 2011). In this experiment growth stimulators promoted seed germination as a result of early initiation and improved stress tolerance by increasing the build-up of enzymes and antioxidant activities under stress condition (Chen and Arora, 2013). Again, growth stimulator promoted transcription factor of isoezymes and gene expression and antioxidants, carbohydrate metabolism and cell development and response to oxidative stress under stress condition (Chen and Arora, 2013). Many studies had demonstrated that pretreatment with ascorbic acid facilitates the retention stay green qualities, primarily leading to increased leaf area trapping sunlight (Keskitalo *et al*., 2005).The experiment proves this truth, ascorbic acid aid plant to do better in any adverse conditions (Torres et al., 2006).

## Conclusion

This present study clearly demonstrated that growth stimulators have a fundamental impact in the initiation of various growth parameters in Sorghum. The plant growth stimulators could be successfully harnessed for enhancement of seed yield, directly or indirectly, through its component.

## REFERENCES

Abdel-Latef, A.A.M (2003). Response of some sorghum cultivars to salt and hormonal treatment. Master Science. Thesis, Faculty of Agriculture South Valley.Qena, Egypt.

Abdel-Latef, A.A., Shaddad, M.A.K.,Ismail, A.M., AbouAlhamed, M.F, (2009) Benzylasemine can alleviate saline injury of two roselle (Hibiscus sabdariffa) cultivars via equilibration of cytosolutes including anthocyanins. International Journal of Agricultural Biology 11:151–157.

Abdul, K.S., Saleh, M.M.S and Omer, S.J (1988). Effects of gibberellic acid angcycocel on the growth, flowering and fruiting characteristics of pepper. Irai Journal of Science6: 7–18.

Abou Al-hamd, M.F (2007). The interactive effects of salinity and phytohormones on some physiological studies of two Hiboiscussabdriffa cultivars. Master Science. Thesis, Faculty Science. South Valley University Qena, Egypt.

Adnan, M., Zahir, S., Fahad, S., Muhammad, A., Mukhtar, A., and Imtiaz, A.(2018). Phosphate solubilizing bacteria nullify the antagonistic effect of soil calcification on bioavailability of phosphorus in alkaline soils. Science.Report 7: 1613–1623.

Ali, H.M., Siddiqui, M.H., Basalah, M.O., Al-Whaibi, M.H., Sakram, A.M. and Al-Amri, A (2012). Effects of gibberellic acid on Growth and photosynthetic pigments of Hibiscussabdariffa L. under salt stress. African Journal of Biotechnology 11(4): 800–804.

Afzal, I., Munir, F., Ayub, C.M., Basra, S.M.A., Hameed, A., Nawaz, A(2009). Changes in antioxidant enzymes, germination capacity and vigour of tomato seeds in response of priming with polyamines. Seed Scienceand Technology 37:765–770.

Afroz, S., Mohammed, F., Hayat, S., and Siddiqui, M.H(2005). Exogenous application of gibberellic acid counteracts the effect of sodium chloride in mustard. TurkJournal of Biology 29:233–236.

Akter, A., Ali, E., Islam., M.M.Z., Karim, R. and Razzaque, A.H.M (2007). Effect of GA3 on growth and yield of mustard. International Journal of Sustainable Crop Production 2: 16–20.

APHA (2008). Standard Methods for the Examination of Water and Wastewater. American Public Health Association, Washington, D.C. USA. 874p.

Ashraf, M., and Rauf, H. (2001). Inducing salt tolerance in maize (Zeamays L.) through seed priming with chloride salts: Growth and ion transport at early growth stages. ActaPhysiologiaePlantarum. 23: 407–414.

Ashraf, M.Y., Azhar, N., and Hussain, M (2006). Indole acetic acid (IAA) induced changes in growth, relative water contents and gas exchange attributes of barley (Hordeumvulgare L.) grown under water stress conditions. Plant Growth Regulation 50:85–90.

Ahmad, I., Khaliq, T., Ahmad, A., Basra, S.M.A., Hasnain, Z and Ali, A (2012) Effects of seed priming with ascorbic, salicylic acid and hydrogen peroxide on the emergence vigourandantioxidant activities of maize. African Journal of Biotechnology 11: 1127–1137.

Anumalla, M., Mallikarjuna, B., Annamalai. A and Jauhar, A(2019). Tolerance of iron deficient and toxic soil conditions in rice. Plants 8(2): 31–39.

Akter, A., Ali, E., Islam, M.M.Z., Karim, R and Razzaque, A.H.M (2007). Effect of GA3 on growth and yield of mustard. International Journal of Sustainable Crop Production 2(2): 16–20.

Angadi, S. V and Entz, M. H (2002). Water relations of standard height and dwarf sunflower cultivars. Crop Science 42: 152–159.

Assefa, M. K., Hunje, R and Koti, R. V (2010). Enhancement of seed quality in soybean following priming treatment. Karnataka Journal of Agricultural Science 23:787–789.

Arora, A., Sairam, R.K and Srivastava, G.C (2002). Oxidative stress and antioxidative system in plants. Curriculum of Science 82: 1227–1338.

Baki, A.A and Andersen, J.D (1972). Physiological and biological deterioration of seeds. In:Seed Biology, Academic Press, New York vol.11.

Bailly, C., Benamar, A., Corbineau, F and Come, D (2000). Antioxidant systems in sunflower (Helianthus annuusL.) seeds as affected by priming. Seed Science Research 10:35–42.

Bradford, K. J (1986). Manipulation of seed water relations via osmotic priming to improve germination under stress conditions. Horticultural Science 21: 1105–1112.

Bassi, G., Sharma, S., and Gill, B. S (2011). Pre-sowing seed treatment and quality invigouration in soybean {Glycinemax (L) Merrill}. Seed Research 31: 81–84.

Bode, K., Doring, O., Luthje, S., Neue, H.U and Bottger, M (1995). The role of active oxygen in iron tolerance of rice (Oryzasauva L.). Protoplasma 184: 249–255.

Bora, R.K., and Sarma, C.M (2006). Effect of gibberellic acid and cycocel on growth, yield and protein content of pea. Asian Journal of Plant Sciences 5: 324–330.

Cavusoglu, K., and Billir, G (2015). Effects of ascorbic acid on the seed germination.Seedling growth and leaf anatomy of barley under salt stress. Journal of Agricultural and Biological Science 10(4):123–129.

Caro, AandPuntarulo (1996). Effect of in vivo iron supplement on oxygenradical production by soybean roots. Biochemistry Biophysic.Acta 1291: 245–251.

Cho, D.Y, and Ponnamperuma, F.N,(1971). Chemistry of submerged soils. Soil Science 112: 184–194.

Clayton, W.D and Renovoize, S.A (1986). Genera Graminum: Grasses of the World, Kew Bulletin Additional Series, Vol. 13, Royal Botanic Gardens, London.

Dayou, E.D., Rakoto, B and Zokpodo, B.L (2017). Physical behaviour of andosols under different levels of mechanization: case of malagasy highlands. In:International Research Journal of India. 2(7):11.

Dorlodot, S., Lutts Sand Bertin, P (2005). Effects of ferrous iron toxicity on the growth and mineral composition of and interspecific rice. Journal of Plant Nutrition 28: 1–20.

Emogor, V (2007). Gibberellic acid (GA3) influence on vegetative growth, nodulation and yield of cowpea (Vignaunquiculata L.). Walp Journal of Agronomy 6:509–517.

Espanany, A., Fallah, S and Tadayyon, A (2016). Seed priming improves seed germination and reduces oxidative stress in black cumin (Nigella sativa) in presence of cadmium. Industrial Crop Production. 79:195–204.

FAO, (1995). Sorghum and Millets in Human Nutrition, FAO Food and Nutrition Series, No. 27, Food and Agriculture Organization, Rome.

Garber, E.D (1950). “Cytotaxonomic studies in the genus Sorghum”, University of California Publications in Botany, 23: 283–361.

Gyaneshwar, P., Kumar, G., Parekh, L and Poole, P (2002). Role of soil microorganisms in improving P nutrition of plant. Plant Soil Journal 245:83–93.

Gamboa-de Buen, A., Cruz-Ortega, R., Martinez-Barajas, E., Sanchez-Coronado, M.E and Orozco-Segovia, A(2006). Natural priming as an important metabolic event in the life history of Wigandiaurens(Hydrophyllaceae)seeds. Physiology of Plant.128:520–530.

Hameed, A.,Bibi, N., Akhter, J and Iqbal, N. (2011). Differential changes in antioxidants, proteases, and lipidperoxidation in flag leaves of wheat genotypes under different levels of water deficit conditions. Plant Physiology and Biochemistry 49:178–185.

Hameed, A., Goher, M and Iqbal, N (2012). Heat stress-induced cell death, changes in antioxidants, lipidperoxidation, and protease activity in wheat leaves. Journal of Plant Growth Regulation 31:283–291.

Harris, D., Rashid, A., Hollington, A., Jasi, L and Riches, C (2007). Prospects of improving maize yield with on farm seed priming. In:Rajbhandari, N.P and Ransom, J.K. ‘Sustainable Maize Production Systems for Nepal’. NARC and CIMMYT, Kathmandu, pp 180–185.

Harlan, J.R and Stemler, A(1976). Plant domestication and indigenous African agriculture. In Harlan, J.R., DE Wet, M.J.J. and Stemler, A. (eds). Origins of African plant Domestication. Mouton, TheHagne, The Netherlands. pp 465–471.

Harrington, J. F., Rappaport L and Hood, K. J (1996). The influence of gibberellins on stemelongation and flowering on endive. Sci. 125: 601.

Hernandex, P (1997). Morphogenesis in sunflower as effected by exogenous application of plant growth regulators. Agriscientia 13:3–11.

Heydecker, W (1973). Accelerated germination by osmotic seed treatment. Nature 246: 42–44.

Hussain, K., Hussain, M., Majeed, A., Nawaz, K., Nisar, M. F and Afghan, S (2010) Morphological response of scurf pea (PsoraleacorylifoliaL.) to indole acetic acid (IAA) and nitrogen (N). World Applied Sciences Journal 8: 1220–1225.

Imasuen, O.I and Onyeobi, T. U. S (2013). Chemical compositions of soils in parts of Edo State, Southwest Nigeria and their relationship to soil productivity. Journal of Applied Science Environmental Management 17: 379–386.

Ikhile, C.I (2016). Geomorphology and hydrology of the Benin region, Edo state, Nigeria. International Journal of Geoscience 7: 144–157.

Ikhajiagbe, B., (2016). Possible adaptive growth responses of Chromolaenaodorata during heavy metal remediation. Ife Journal of Science 18: 403–411.

Ikhajiagbe, B., Musa, S.I and Okeme, J.O (2019). Effect of changes in soil cation exchange capacity on the reclamation of lead by Eleusineindica (L) Gaertn. FUDMA Journal of Sciences 3(4): 176–183.

Ishibashi, Y and Iwaya-Inoue, M (2006). Ascorbic acid suppresses germination and dynamic states of water in wheat seeds. Plant Production Science. 9(2): 172–175.

Joshis, X., Cho, C., Racz Gand Chang, C (2007). Chemical retardation of phosphate diffusion in an acid soil as affected by liming. Nutrient Cycle Agroecosystem 64: 213–224.

Kampfenkel Kand Montagu, V (1995). Effects of iron excess on Nicotianaplumbaginifoliaplants (implications to oxidative stress). Plant Physiology 107: 725–735

Khan, T.A., Mazid Mand Mohammad, F (2011). A review of ascorbic acid potentialities against oxidative stress induced in plants. Journal of Agrobiology 28(2):97–111.

Khan, H. A., Ayub, C. M., Pervez, M. A., Bilal, R. M., Shahid, M. A and Ziaf, K (2009). Effect of seed priming with NaCl on salinity tolerance of hot pepper (Capsicum annuum L.) at seedling stage. Soil Environment 28: 81–87.

Kaur, S., Gupta, A.K and Kaur, N (2000). Effect of GA kinetin and indole acetic acid on carbohydrate metabolism in chickpea seedling germinating under water stress. Plant Growth Regulation 30: 61–70.

Min, L., Li, Y., Hu, Q., Zhu., L., Gao, W., Wu, Y., Ding, Y., Liu, S., Yang, X and Zhang, X (2014). Sugar and auxin signalling pathways respond to high-temperature stress duringantherdevelopment as revealed by transcript profiling analysis in cotton. Plant Physiology 164:1293

Mwale, S. S., Hamusimbi, C and Mwansa, K (2003). Germination, emergence and growth of sunflower (Helianthus annuus L.) in response to osmotic seed priming. Seed Science Technology 31: 199–206.

Nnadia, E.O., Mbah, C.N., Nweke, A and Njoku, C(2019). Physicochemical properties of an acid ultisol subjected to different tillage practices and wood-ash amendment: Impact on heavy metal concentrations in soil and Castor plant. Soil and Tillage Research 194: 104–288.

Peng, X.X and Yamauchi, M (1993). Ethylene production in rice bronzing leaves induced by ferrous iron. Plant Soil 149: 227–234.

Rahimi, A. (2013) seed priming improves the germination performance of cumin (CuminumcyminumL.) under temperature and water stress. Industrial Crop Production 42: 454–460.

Raza, S.H., Shafiq, F and Chaudhary, M (2013). Seed invigoration with water, ascorbic and salicylic acid stimulates development andbiochemical characters of okra (Abelmoschusesculentus) under normal and saline conditions. International Journal of Agriculture and Biology 15(3): 486–492.

Sadeghian, S.Y and Yavari, N (2004). Effect of water deficient stress on germination and early seedling growth in sugar beet. Journal of Agronomy Crop Science 190: 138–144.

Salehzade, H., Shishvan, M. I., Ghiyasi, M., Forouzin, F. and Siyahjani, A. A. (2009). Effect of seed priming on germination and seedling growth of wheat [Triticumaestivum(L).]. Research Journal of Biology Science 4: 629–631.

Seyyedi, S.M., Khajeh-Hosseini, M., Moghaddam, P.R and Shahandeh, H (2015). Effects of phosphorus and seed priming on seed vigor, fatty acids composition and heterotrophic seedling growth of black seed (NigellasativaL.) growth in calcareous soil. Industrial Crop Production 74: 939–949.

Snowden, R.E.D and Wheeler, B.D (1995). Iron toxicity of fen plant species. Journal of Ecology 81:35–46.

Stefoska-Needham, A., Beck, E.J., Johnson, S.K and Tapsell, L.C (2015). Sorghum: an underutilized cereal whole grain with the potential to assist in the prevention of chronic disease. Food Reviews International 31(4): 401–437.

Taiz, L and Zeiger, E (2010). Plant physiology, 3rd edition, Sinauer Associates, Inc., Publishers Sunderland Masschusetts USA. p.690.

Talla, S., Riazunnisa, K., Padmavathi, L., Sunil, B., Rajsheel, P and Raghavendra, A.S. (2011). Ascorbic acid is a key participant during the interaction between chloroplasts and mitochondria to optimize photosynthesis and protect against photoinhibition. Journal of Biosciences 36(1): 163–173.

Tavanoglu, C., Catar, S, Sand Ozudogru, B (2015). Fire related germination and early seedling growthin herbaceous species on central Anatolian steppe. Journal of Arid Environments 122: 109–116.

Taylor, A. G., Allen, P. S., Bennett, M. A., Bradford, K. J., Burrisand, J. S and Misra, M. K (1998). Seed enhancements. Seed Science Research 8: 245–256.

Valera, L (1977). Phosphorus availability and sorption as affected by long-term fertilization. Agronomy Journal 106: 1584–1592.

Wang, Y. L., Tang, J. W., Zhang, H. L., Schroder, J. L and He, Y. Q (2014). Phosphorus availability and sorption as affected by long-term fertilization. Agronomy Journal 106: 1584–1592.

Wang, H.Y., Chen, C.L and Sung, J.M (2003). Both warm water soaking and solid priming treatments enhanceanti-oxidation of bitter gourd seeds germinated at sub-optimal temperature. Seed Science Technology 31:47–56.

Wiersema, J.H and Dahlberg, J (2007). “The nomenclature of Sorghumbicolor (L.) Moench (Gramineae)”, Taxon. 56: 941–946.

Youssef, A.S.M. (2004). Physiological studies on growth and flowering of Sterilitiziareginaeplant. Ph.D.Thesis Faculty of Agriculture MoshtohorZag University.

Yu, H., Li, B., Liu, S., Huang, W., Liu, T and Yu, W(2016). Iron redox cycling coupled to transformation and immobilization of heavy metals: Implications for paddy rice safety in the red soil of south China. Advances in Agronomy 137: 65–73.

Zhang, Y (2013) ascorbic acid in plants: biosynthesis, regulation and enhancement. 1a ed. New York: Springer-Verlag p117.

Zhang, M., Zheng, X., Song, S., Zheng, Q., Hou, L., Li, D., Zhao, J., Wei, Y., Li, X., Luo, M., Xioao, Y., Luo, X., Zhang, J., Xiang Cand Pei, Y (2011). Spatiotemporal manipulation of auxin biosynthesis in cotton ovule epidermal cells enhances fiber yield and quality. National Biotechnology 29: 453–458.

